# Deconvolution of ears’ activity (DEA): A new experimental paradigm to investigate central auditory processing

**DOI:** 10.1101/2020.07.11.198929

**Authors:** Fabrice Bardy

## Abstract

A novel experimental paradigm, “deconvolution of ears’ activity” (DEA), is presented which allows to disentangle overlapping neural activity from both auditory cortices when two auditory stimuli are presented closely together in time in each ear.

Pairs of multi-tone complexes were presented either binaurally, or sequentially by alternating presentation order in each ear (i.e., first tone complex of the pair presented to one ear and second tone complex to the other ear), using stimulus onset asynchronies (SOAs) shorter than the neural response length. This timing strategy creates overlapping responses, which can be mathematically separated using least-squares deconvolution.

The DEA paradigm allowed the evaluation of the neural representation in the auditory cortex of responses to stimuli presented at syllabic rates (i.e., SOAs between 120 and 260 ms). Analysis of the neuromagnetic responses in each cortex offered a sensitive technique to study hemispheric lateralization, ear representation (right versus left), pathway advantage (contra- versus ipsi-lateral) and cortical binaural interaction.

To provide a proof-of-concept of the DEA paradigm, data was recorded from three normal-hearing adults. Results showed good test-retest reliability, and indicated that the difference score between hemispheres can potentially be used to assess central auditory processing. This suggests that the method could be a potentially valuable tool for generating an objective “auditory profile” by assessing individual fine-grained auditory processing using a non-invasive recording method.

## Introduction

The auditory system is a binaural system. Auditory cortices in right and left hemispheres receive ascending projections originating from each ear. The resulting activity in one cortex is a mixture of signals from both ears. The effects of monaural and binaural stimulation on cortical responses have been studied considerably in humans, using techniques such as magnetoencephalography (MEG) [Pantev et al., 1986]. MEG is well suited to study hemispheric processing differences given the low dispersion of the magnetic field and the location of the cerebral auditory cortical centers in the temporal lobe of each hemisphere. For monaural sound presentation, there is evidence of a predominant contra-lateral pathway in the human auditory system [Mäkelä et al., 1993; Pantev et al., 1986; Pantev et al., 1998]. The contra-lateral advantage is characterized by shorter latencies and larger amplitudes of the N100m. These measures reflect anatomical differences, especially the larger number of neurons projecting on the contra-lateral compared to the ipsi-lateral side of the ascending auditory pathways. For binaural presentation at the cortical level, MEG frequency-tagging of cortical steady-state responses can be employed [Fujiki et al., 2002]. Here, stimuli receive a marker, or tag, using a specific modulation frequency. This makes it possible to identify which stimulus evoked the observed cortical response.

The auditory system is a temporally fast system. It can process acoustic stimuli presented with short temporal disparities between the ears. Processing rapidly changing sounds encompasses several levels of transformation from one cochlea to the auditory cortex of both hemispheres. Unfortunately, a non-invasive objective measure of binaural interaction in the auditory cortex during rapid stimulation with temporally restricted sounds is not yet available. However, if such a method were to be available, research on the interaction and/or integration of signals in the auditory cortex for stimuli presented at syllabic rates (i.e., between 4 and 10 Hz) could provide new insights into normally developed and disordered central auditory processing systems.

This report describes a novel experimental paradigm, named “deconvolution of ears’ activity” (DEA), which makes use of the least-squares (LS) deconvolution technique to allow separation of left and right ear activity in each hemisphere to rapidly presented stimuli [Bardy et al., 2014a; Bardy et al., 2014b].

The LS deconvolution technique is a mathematical algorithm designed to disentangle temporally overlapping brain responses. The technique relies on the timing characteristics of the stimulus sequence to be unequally spaced. This specific property is called ‘jitter’. In the DEA paradigm, LS deconvolution is applied to a sequence of stimuli presented in pairs either binaurally or sequentially, using stimulus onset asynchronies (SOAs) shorter than the duration of the cortical response. Right and left ear activity is extracted from the mixture of signals in both auditory cortices such that, using this method, the signal propagation from each ear to each auditory cortex can be tracked. The DEA paradigm is introduced in this paper, and is evaluated on three normal hearing adults as a proof-of-concept.

Two hypotheses were investigated: (1) the LS deconvolution technique can disentangle temporally overlapping brain responses in each auditory cortex originating from both ears with a high test-retest reliability; and (2) an auditory profile can be generated based on measures of the auditory pathway lateralization, hemispheric advantage, ear advantage and binaural cortical interaction.

## Methods

Subjects. Test and retest MEG data were obtained from 3 right-handed adult subjects (3 males, age: 37, 32, 29) on two separate occasions. Subjects had no history of neurological or audiological problems and had pure tone audiometric thresholds less than or equal to 20 dB HL in all octave frequencies between 250 to 8000 Hz. This study was approved by and conducted under oversight of the Macquarie University Human Research Ethics Committee. All subjects gave written informed consent to participate in this study.

Stimulation. Two multi-tone (MT) stimuli, selected to optimize the amplitude of the cortical response [Bardy et al., 2015], were obtained by amplitude-modulated tone-bursts composed of carrier frequencies of 2 and 1 kHz with modulation frequencies 800 and 400 Hz respectively. The two MTs were presented in pairs, using jittered SOAs with means of 120, 190 or 260 ms. The jitter distribution, permitting the deconvolution, was rectangular with a width of 70 ms and a step size of 13.3 ms. The inter-pair interval (IPI), representing the time interval between the onset of two successive pairs of stimuli, was jittered with 400 ms around an average of 1400 ms. The MTs had a rise and decay time of 10 ms, a duration of 50 ms and an rms intensity of 70 dB SPL. They were presented through shielded transducers [Oldfield, 1971]. The stimuli were presented in 3 presentation conditions. The first presentation condition was binaural (both stimuli of the pair presented simultaneously to the right and left ears). In the two other presentation conditions, stimuli were alternated sequentially in each ear (i.e., when the left ear received the first tone, the right ear received the second tone of the pair, and vice versa). All 9 conditions (3 SOAs x 3 presentation conditions) were randomly presented in a 25 minute long stimulus sequence. In conditions where the cortical response was longer than the SOA, brain responses overlapped in time, and LS deconvolution described by Bardy et al. [2014a] was employed to disentangle the occurring overlapping responses. Thus, for example, in the alternating sequential condition, it was possible within each auditory cortex to separate the activity elicited by the stimulus to the right and left ears respectively from the overlapping cortical response (Figure 1).

**Figure 1.**
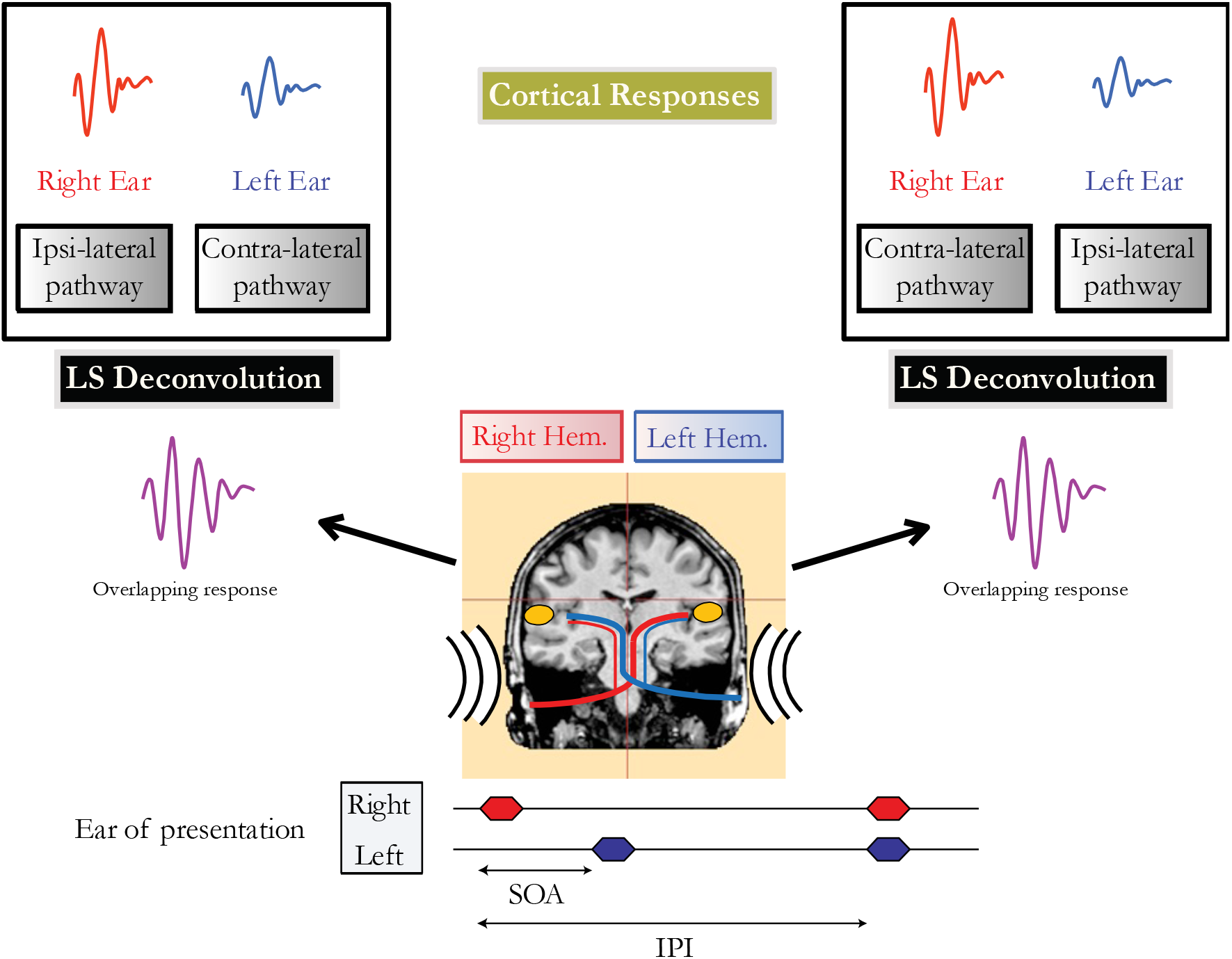
Schematic representation of a sequential condition. The first auditory stimulus of the pair is presented to the right ear, followed closely by a presentation to the left ear. The overlapping neural activity reaches both hemispheres via contra- and ipsi-lateral pathways. The overlapping responses from both ears in each auditory cortex are unwrapped using the least squares (LS) deconvolution technique.

Procedure. MEG data were continuously recorded using a whole-head MEG system (Model PQ1160R-N2, KIT, Kanazawa, Japan) consisting of 160 coaxial first-order gradiometers with a 50 mm baseline [Kado et al., 1999; Uehara et al., 2003]. MEG data were acquired in a magnetically shielded room using a sampling rate of 1000 Hz with a bandpass filter of 0.1–200 Hz and a 50 Hz notch filter. For co-registration, the location of five indicator coils placed on the participant’s head were digitized. A pen digitizer (Polhemus Fastrack, Colchester, VT) was used to measure the shape of each participant’s head which was then carefully centered in the MEG dewar (position error <10 mm for each subject). Artefact removal from MEG data included signals exceeding amplitude (>2700 fT/cm) and magnetic gradient (>800 fT/cm/sample) criteria [Yetkin et al., 2004]. Averaging and band-pass filtering between 3 Hz (6 dB/octave, forward) and 30 Hz (48 dB/octave, zero-phase) was performed for each trigger condition using the non-contaminated epochs. The accepted epochs after artefact rejection were exported from BESA 5.3 into MATLAB (MathWorks, Natick, MA) and downsampled to 100 Hz. Deconvolution was performed for each of the 160 channels to disentangle overlapping responses. For each condition, recovered responses were defined by epochs of 100 ms pre-stimulus to 380 ms post-stimulus.

Statistical analysis. Amplitudes and latencies were defined by peak measures of magnetic global field power (mGFP) calculated on 40 sensors located over the temporal lobe in each hemisphere. For each subject and each condition, the N100m was defined as the most positive peak in the 80-150 ms following the sound onset. The selected time window for the P200m was 120-200 ms. Two repeated measures ANOVAs were performed. Greenhouse-Geisser corrections for sphericity were applied, as indicated by the the cited ε value.

Individual laterality indices (LIs) for hemisphere, pathway, ear and cortical binaural interaction were calculated. For each subject, LIs were calculated based on the relevant mGFP response amplitudes, time-averaged over a 200-ms window post-onset. Figure 2 displays an example of auditory cortical responses elicited by pairs of auditory stimuli presented binaurally or alternated sequentially for an individual subject with SOAs jittered around 190 ms. For hemispheric lateralization, the LI was calculated as the difference between left and right mGFP response amplitudes (bottom versus top 6 panels in Figure 2) normalized by the sum of left and right mGFP responses. The LI was +1 for a response geared completely asymmetrical towards the left hemisphere, zero for a symmetrical response, and −1 for a response geared completely asymmetrical towards the right hemisphere. For pathway advantage, the LI was calculated employing the same method using the responses associated with the contra- (panels labeled 3R, 4L, 5L and 6R in Figure 2) and the ipsi-lateral pathways (panels labeled 3L, 4R, 5R, 6L in Figure 2). The ear LI was calculated by comparing mGFP responses from the left ear (3^rd^ and 6^th^ columns in Figure 2) to the responses from the right ear (4^th^ and 5^th^ columns in Figure 2). Finally, the binaural interaction LI was computed by comparing binaural stimulation (first 2 columns in Figure 2) and monaural stimulation responses (last 4 columns in Figure 2). The binaural interaction LI was computed for both hemispheres and for each pathway (i.e., ipsi- and contra-lateral). For each subject, the difference between the means for each LI was checked by the Student’s t-test. The threshold for significance after Bonferroni correction was p<0.0041. Test-retest reliability indices were obtained using the mean squared error for each measure of LI as well as the intra-class correlation coefficients (ICCs) on mGFP waveforms.

**Figure 2.**
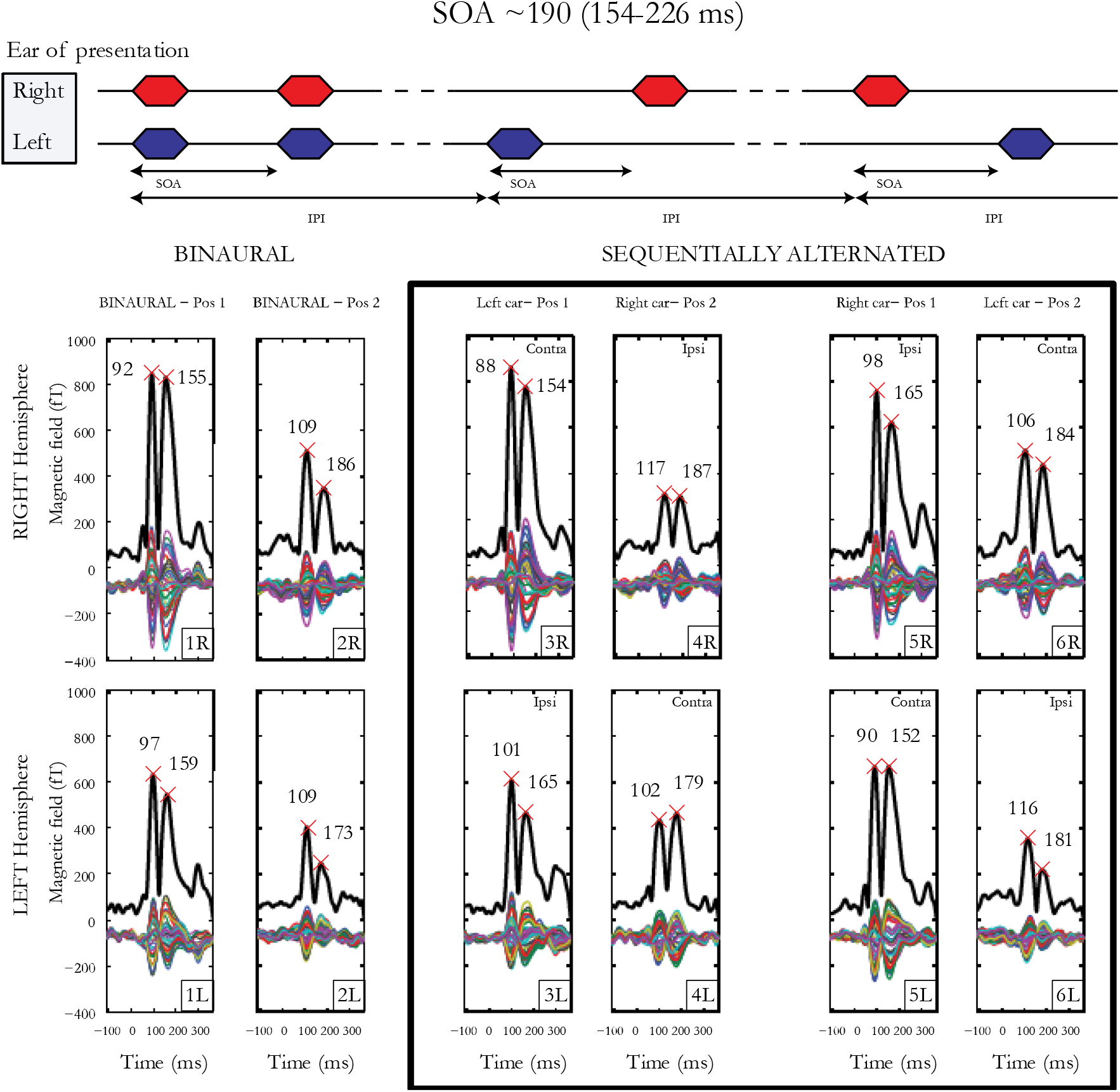
Cortical responses from subject 1 for SOAs jittered around 190 ms. Multiple thin waveforms represent activity recorded by each of the 40 sensors located over the temporal lobe, in each hemisphere, after LS deconvolution, from −100 to 380 ms after stimulus onset. mGFP waveforms are represented with a thick black line, provided for both right and left hemispheres, the 3 presentation conditions (1 × binaural, 2 × sequentially alternated) and both first and second tone-bursts. Latencies of the N100m and P200m are indicated by crosses.

## Results

### Cortical responses to rapidly presented stimuli

Figure 3 presents means and standard deviations of N100m and P200m amplitudes and latencies for ear, stimulus, pathway, and hemisphere. The first ANOVA with hemisphere (right, left), presentation condition (binaural, sequentially alternated left ear first, sequentially alternated right ear first), SOA (~120, ~190, ~260 ms), and position (first or second stimulus in the pair) as factors, revealed that the response to the second stimulus of the pair was significantly smaller in amplitude and longer in latency compared to the response to the first stimulus of the pair for both N100m (Amp. F(1,5)=77.55, p=0.0003, ɛ = 1; F(1,5)=524.59; Lat. p=0.000003, ɛ = 1) and P200m (Amp. F(1,5)=106.95, p=0.0001, ɛ = 1; Lat. F(1,5)=41.92, p=0.001, ɛ = 1).

**Figure 3.**
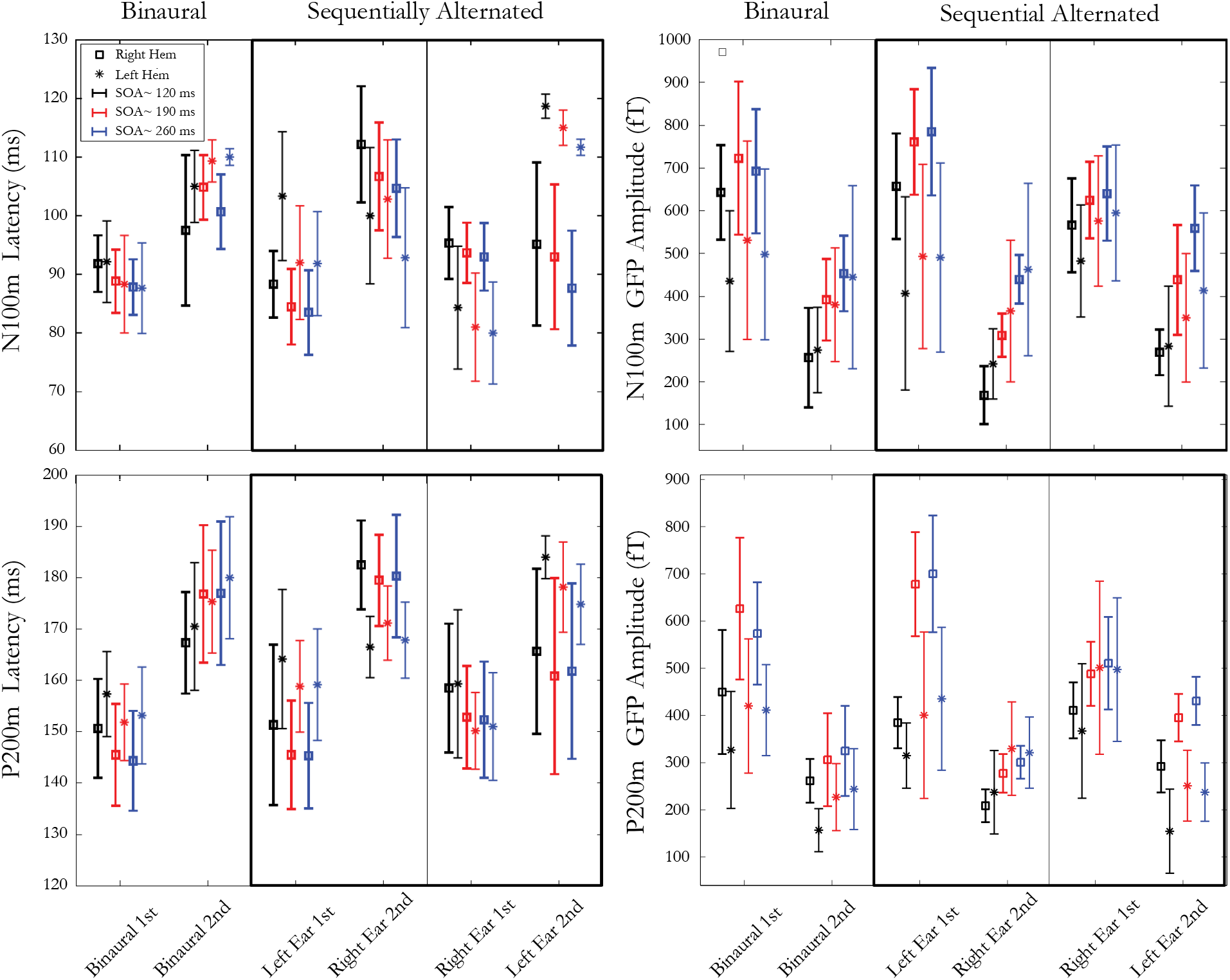
Latencies (left panels) and amplitudes (right panels) of mGFP N100m (top panels) and P200m (bottom panels) components. Each panel represents 3 presentation conditions: 1 binaural condition and 2 sequentially alternated conditions (stimulus presented first at either left or right ear). Within each presentation condition, three SOAs (~120, ~190 and ~ 260 ms) are used, resulting in two responses to both stimuli of the pair, recorded from both right and left hemispheres. Error bars denote standard deviations between participants.

The second ANOVA similarly used hemisphere, presentation condition and SOA as factors, but only considered responses to the second stimulus in the pair. The effect of SOA was found to be significant for both amplitudes and latencies of N100m (Amp. F(2,10)=46.48, p=0.000009, ɛ = 0.58; Lat. F(2,10)=7.30, p=0.03, ɛ = 0.54) and for P200m amplitude (F(2,10)=53.95, p=0.000004, ɛ = 0.78). The cortical responses to the second stimulus in the pair increased in amplitude and decreased in latency for longer SOAs when compared to shorter SOAs. Moreover a significant interaction was present between SOA and presentation condition for both N100m (F(4,20)=10.07; p=0.001, ɛ = 0.60) and P200m (F(4,20)=8.29; p=0.004, ɛ = 0.60) latencies. A decrease in response latency was observed when SOA increased in the sequentially alternated presentation condition, while this trend was inverted in the binaural presentation conditions.

### Hemispheric lateralization

The hemispheric lateralization index (LI) for response amplitude presented in Figure 4a shows intra-individual differences on the vertical abscissa, and inter-individual differences on the horizontal abscissa. Subject 1 presented a rightward, subject 2 a large rightward, and subject 3 a slightly leftward lateralization. The t-test, which allows comparing the hemispheric LI to 0, was significant for each subject (p<0.001) after Bonferroni correction. No differences in symmetrical activation were found for the latencies either for subject 1 (p=0.86) or subject 2 (p=0.51). However, significantly earlier latencies were found in the right hemisphere for subject 3 (p=0.0003).

**Figure 4.**
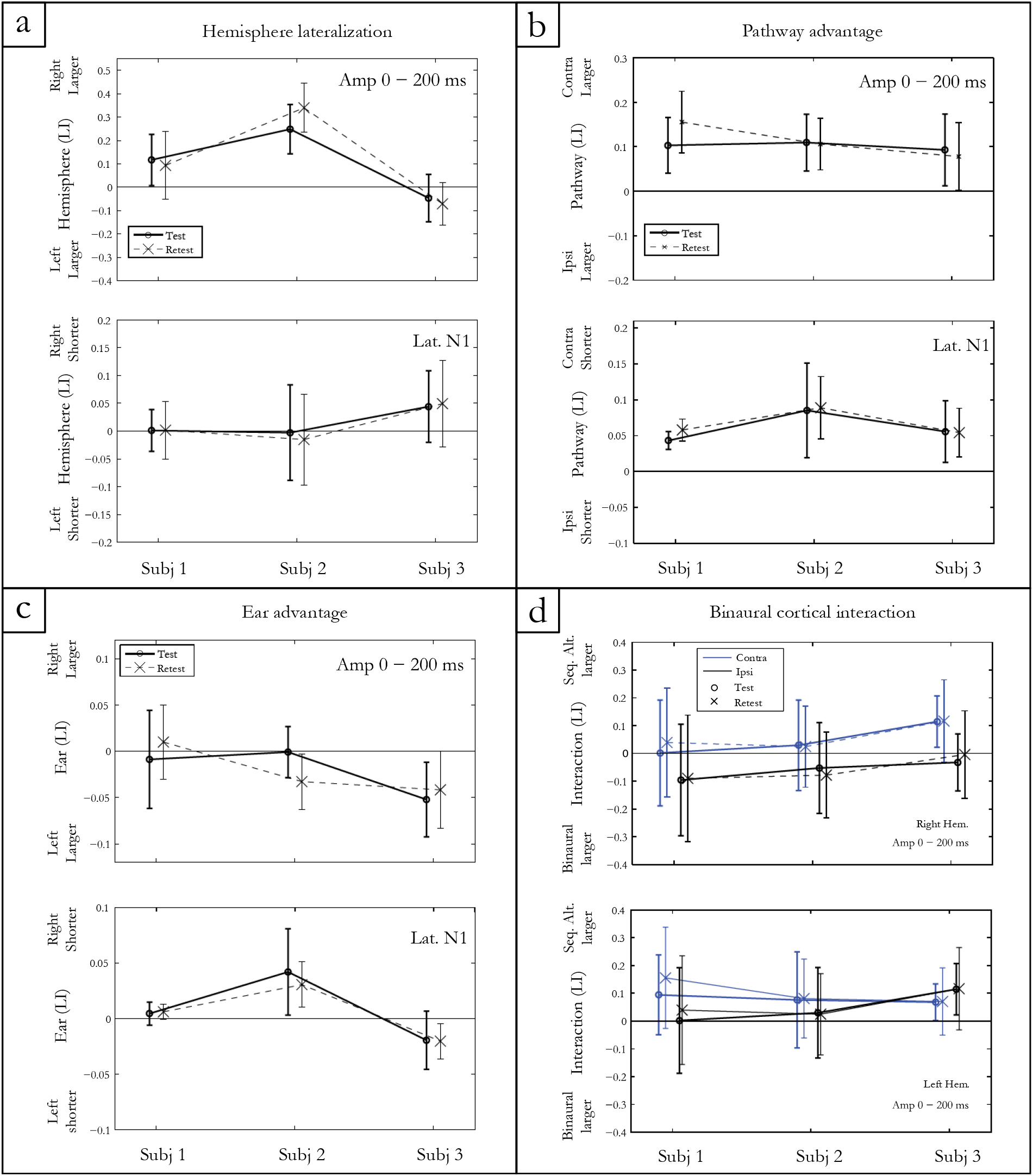
Indices of hemispheric lateralization, pathway advantage, and ear advantage for mGFP amplitudes in a 200 ms post onset window, and for N100m latency. Both test and retest conditions are shown. The binaural cortical interaction is represented for the mean mGFP amplitude in the right and left hemisphere for the contra- and ipsi-lateral pathways. Error bars denote standard deviations between conditions for each participant.

### Pathway advantage

The pathway LI calculated by contrasting contra- versus ipsi-lateral pathway responses in the sequential conditions is represented in Figure 4b. After Bonferroni correction, significantly larger amplitudes and shorter latencies for the N100m and P200m were measured in the contra-lateral pathway for all subjects (p<0.0001).

### Ear advantage

The statistical results of ear LI presented in Figure 4c indicated no significant amplitude difference between the activity elicited by the right and the left ear for subject 1 (p=0.96) and for subject 2 (p=0.01). A left ear advantage was observed for subject 3 for both amplitude (p=0.002) and latency (p=0.002).

### Cortical binaural interaction (CBI)

Figure 4d shows the CBI for the 3 subjects in both hemispheres for contra- and ipsi-lateral pathways. The finding of a positive CBI LI indicates that the response recorded in the sequentially alternated condition is larger compared to the response in the ipsi-lateral pathway. CBI of different natures are observed for each subject. When collapsed across hemispheres, the t-test showed that CBI was close to significance only for subject 3 (subject 1: p = 0.02; subject 2: p =0.10; subject 3: p=0.006).

### Test-retest reliability

Two different test-retest reliability measures were computed. First, the mGFP waveforms were compared for test and retest conditions by computing the intra-class correlation coefficients (ICCs) for the 3 subjects in a 250 ms window post onset. A mean ICC value larger than 0.75 for each subject (i.e., subject 1 = 0.78, subject 2 = 0.79; subject 3 = 0.84) demonstrated a good test-retest reliability.

Second, a test-retest index was calculated using the mean squared error (mean = 0.057; SD = 0.026) of all four indices presented in Figure 4 (i.e. hemispheric lateralization, pathway advantage, ear advantage and CBI).

## Discussion

The central aim of this paper was to introduce the deconvolution of ears’ activity (DEA) paradigm which disentangles the activity in both auditory cortices elicited by stimuli presented to both ears simultaneously or separately. In this paradigm, the LS deconvolution technique was applied to MEG data recorded using pairs of stimuli presented either binaurally or alternating sequentially (i.e. right-left and left-right). The DEA paradigm allowed the investigation of auditory information transfer from one specific ear to both auditory cortices. It could also be used to explore response lateralization, the strength of crossed auditory pathways and the response adaptation properties to auditory stimuli closely separated in time. Furthermore, it allowed for the investigation of non-linear processing in the brain and CBI, mainly caused by inhibition mechanisms [Imig and Brugge, 1978; Imig and Reale, 1981; Papanicolaou et al., 1990; Reite et al., 1981].

We demonstrated the feasibility and test – retest reproducibility of this non-invasive measure on 3 right-handed normal-hearing subjects. The case studies provided examples of different auditory processing characteristics at the cortical level, identifiable at the individual level. The inter-individual differences were detectable by assessment of the difference in response between experimental conditions. For example, hemispheric lateralization was assessed by computation of the LI calculated from the responses in each hemisphere. The CBI was investigated by contrasting binaural and monaural stimulation both in contra- and ipsi-lateral pathways. The results collected using the DEA paradigm allows an objective auditory processing characterization and the generation of an individual “auditory profile” in a relatively quick time (i.e., 25 min).

### Experimental results

The data recorded from 3 normal-hearing subjects confirmed that both ears were represented in each cortical hemisphere. However, differences in latency and amplitude were observed for each response to various conditions.

Beyond the idea proposed by Poeppel [2003] that sound processing in the brain is a bilateral phenomenon, the present study revealed inter-individual differences in the hemispheric lateralization of the cortical response. While two subjects showed a rightward hemisphere lateralization for response amplitude, the third subject had a leftward lateralization. These hemispheric asymmetries and specializations for processing auditory stimuli were also reported previously by Mäkelä et al. [1993] and Jamison et al. [2006]. The cerebral lateralization of the auditory cortical area however is still highly debated [Bishop, 2013; Scott and McGettigan, 2013].

For all subjects tested, the N100m was larger and approximately 10 ms shorter for the contra-lateral compared to the ipsi-lateral auditory pathway in the sequentially alternated conditions. These results are in agreement with several studies showing a contra-lateral dominance based on lateralization of the N100m component [Pantev et al., 1986; Pantev et al., 1998; Tiihonen et al., 1989; Woldorff et al., 1999].

Individual differences were also observed when comparing ear activity. Further research will need to investigate whether this objective measure of ear advantage is correlated with behavioral performance on a dichotic listening task such as the Dichotic Digits Test [Musiek, 1983].

The DEA paradigm allowed to investigate the suppression-type interaction and neural mechanisms underlying the processing of rapidly presented signals. As shown in Figure 4d, different binaural interactions were observed. Amplitudes of responses elicited in the sequentially alternated presentation condition were found to be either slightly larger, slightly smaller or of similar amplitude compared to the binaural presentation condition. Inter-subject differences were observed with different interactions depending on hemisphere and pathway involved. A MEG study using complex tones showed that responses to ipsi-lateral stimuli over the right auditory cortex are inhibited by the stimuli presented in the contra-lateral (left) ear [Brancucci et al., 2004].

Lastly, cortical responses to stimulus pairs separated by short SOAs allowed the study of the representation in the auditory cortex of stimuli presented closely together. Results presented in Figure 3 showed a large decrease in amplitude and increase in latency of the cortical response when it stimulus was closely preceded by another auditory stimulus.

We conclude that the DEA paradigm could represent a technique to study interesting properties of the central auditory system. Individual differences are of special interest as they provide an alternative characterization of the hearing profile of a person which could potentially be useful to for example objectively identify auditory processing disorder (APD) subjects. Using the LS deconvolution technique to separate overlapping ear activity in both auditory cortices, recorded MEG data can provide a measure for rapid temporal processing, response lateralization, auditory pathway and ear advantage, and CBI for rapidly presented sound stimuli. Such a test would allow studying the temporal acuity of the human auditory system when processing rapid changes in the acoustic signal. Moreover, it could provide insights concerning the flow of neural signals from the cochlea to the cerebral cortex. From a clinical perspective, tests are needed to better evaluate and understand the neurological characteristics of binaural processing occurring in the auditory system. Such tests could contribute to the diagnosis of APDs or neurodevelopment disorders, such as specific language impairment (SLI) or dyslexia where abnormal crossing pathways or the disability to process rapid auditory stimuli has been identified [Lamminmäki et al., 2012]. More complex sounds, such as speech syllables (using carefully selected jitter parameters), could be used in the future to investigate the influence of stimuli on binaural interaction mechanisms and lateralization of the response.

## Acknowledgments

This work was supported in part by: the HEARing CRC, established and supported under the Australian Cooperative Research Centres Program, an Australian Government Initiative, by the Australian Government Department of Health and by the Oticon Foundation. The authors gratefully thank Bram Van Dun, Harvey Dillon, Catherine McMahon, Robert Cowan and Ramesh Rajan for their suggestions during the preparation of this manuscript.

